# Identification of Thioredoxin1 interacting proteins in neuronal cytoskeletal organization during autophagy

**DOI:** 10.1101/2024.02.27.582366

**Authors:** Md Imamul Islam, Shakila Sultana, Nirmala Padmanabhan, Mahmud-ur Rashid, Tabrez Siddiqui, Kevin Coombs, Peter F. Vitiello, Soheila Karimi-Abdolrezaee, Eftekhar Eftekharpour

## Abstract

Thioredoxin1 (Trx1) is a major cytoplasmic thiol oxidoreductase protein involved in redox signaling. This function is rendered by a rapid electron transfer reaction during which Trx1 reduces its substrate and itself becomes oxidized. In this reaction, Trx1 forms a transient disulfide bond with the substrate which is unstable and therefore identification of Trx1 substrates is technically challenging. This process maintains the cellular proteins in a balanced redox state and ensures cellular homeostasis. Trx1 levels are reduced in some neurodegenerative diseases; therefore, understanding the interactions between Trx1 and its substrates in neurons could have significant therapeutic implications. We utilized a transgenic mouse model expressing a Flag-tagged mutant form of Trx1 that can form stable disulfide bonds with its substrates allowing identification of the Trx1 interacting proteins. The involvement of Trx1 has been suggested in autophagy, we aimed to investigate Trx1 substrate after pharmacologic induction of autophagy in primary hippocampal neurons. Treatment of primary neurons by rapamycin, a standard autophagy inducer, caused significant reduction of neurite outgrowth and alterations in the cytoskeleton. Through immunoprecipitation and mass spectrometry, we have identified 77 Trx1 interacting proteins which were associated with a wide range of cellular functions including a major impact on cytoskeletal organization. The results were confirmed in Trx1 knocked-down cells and in nucleofected primary neurons. Our study suggests a novel role for Trx1 in regulation of neuronal cytoskeleton organization, marking the first investigation of Trx1-interacting proteins in primary neurons and confirming the multifaceted role of Trx1 in physiological and pathological states.

## Introduction

Cysteine is a unique amino acid which contains a reactive sulfhydryl group, and therefore is subject to a variety of post-translational modifications including oxidation, glutathionylation, nitrosylation, and acylation [1–3]. These changes play an important part in determining the protein structure and function. In simple organisms such as, cysteine is found in only 0.4%– 0.5% of proteins, while in more complex organisms such as mammals ∼2.26% of proteins contain cysteine [4]. Cysteine containing proteins play an important role in cellular redox balance, especially in the brain due to its high rate of reactive oxygen/nitrogen species (ROS/RNS). The sulfhydryl group in cysteine containing proteins makes it an important player in redox signaling through reversible oxidation by ROS. Since maintaining cysteine redox status is crucial for neuronal homeostasis, many neurodegenerative diseases are linked to cysteine deficiency [5, 6]. The cells are endowed with a supply of cysteine-containing thiol antioxidants including glutathione (GSH), glutaredoxin, and thioredoxin that have low redox potential and therefore are responsible for scavenging ROS and maintaining the cysteine residue in its reduced form. Mild oxidation of the thiol group on cysteine residue to sulfenic acid (Cys-SOH) is reversible by these antioxidant thiols, further oxidation and formation of sulfinic and sulfonic acids are irreversible, which requires protein degradation by proteasome and autophagy systems [7, 8].

Amongst these thiols, GSH is the major ROS scavenger and is involved in reduction of oxidized protein through protein glutathionylation, resulting in formation of stable protein adducts that can be identified using several chromatography and proteomic approaches [9] and therefore the importance of GSH in redox has been well identified for the cell. The protein adducts will be fully reduced through electron exchange with glutaredoxin [10].

Trx1, participates in regulation of redox balance, has lower ROS scavenging capacity than glutathione and is mostly involved in reduction of disulfide bonds [11]. A growing list of proteins are known to be dependent on Trx1 disulfide reducing capacity, although identification of these proteins is technically challenging due to its mode of action. Trx1 has two cysteines on its active site (Cys32 and Cys35) which catalyze the disulfide bond reduction process [12]. In reduced Trx1, Cys32 first forms an intermolecular disulfide bond by interacting with the oxidized cysteine of the substrate. Immediately, Cys35 of Trx1 attacks the newly formed disulfide bond, causing a formation of an internal disulfide bond between Cys32 and Cys35, and rendering the target proteins/substrates to be reduced [12]. Examples of Trx regulatory roles have been shown under oxidative stress conditions where it directly reduces its substrates such as caspases [13], histone deacetylases (class II) [14], AMP-activated protein kinase (AMPK) [15–17], NLR family pyrin domain containing 1 NLRP1 [18], mechanistic target of rapamycin (mTOR) [19], and nuclear factor κ B NFκB [20] through the above-mentioned thiol-disulfide exchange reactions. Trx1 protein targets are increasingly identified in different systems, however there is currently no information available on Trx1 targets in neurons.

Our lab has previously shown that downregulation of Trx1 is associated with disruption of lysosomal activity and neuronal cell death [21]. Other reports also suggest that Trx1 might be involved in initiation of autophagy through regulation of autophagy related protein-7 (ATG7) in cardiac cells [22], although this remains to be shown in neuronal cells. For this study we used primary cortical neuron cultures from a mouse model in which these cells express a mutant Trx1. The Cys35 in the mutant protein is replaced by a Serine, allowing formation of stable disulfide bonds with the substrate. The mutant protein also contains a Flag-tag which can be used for purification and identification of Thioredoxin-substrate complexes. This approach has been previously used for identification of Trx1 targets in mouse lungs under hyperoxic conditions [23]. We confirmed the induction of autophagy in the neurons after exposure to Rapamycin. This approach revealed novel interactions between Trx1 and neuronal proteins. Amongst those, 77 proteins were differentially expressed between control and those after induction of autophagy. Using pharmacological and genetic inhibition of Trx1 in HT22 cells, we show that Trx1 availability is important for actin fibrilization. This is the first proteomic study examining Trx1 targets in neurons.

## Results

### Rapamycin induces autophagy in primary cortical neurons

Primary cortical neurons were cultured from E17 flag-hTrx1Cys35S expressing mice (Fig. 1A). The primary cortical neurons were treated with 100 nM of rapamycin on DIV 7 for 24 h and cells were harvested. Due to the modification of Cysteine at position 35 with Serine, the flag-tagged Trx1 will be unable to reduce its target and the initial mixed disulfide interacting partners formed by Cys 32 will be trapped. We first examined the expression of endogenous Trx1 (wild type) and the overexpressed flag-hTrx1C35S protein using Anti-Trx1 antibody. Two bands representing the 12kDa endogenous Trx1, and its Flag-tagged hTrx1Cys35S isoforms were detected. The molecular weight difference is due to the presence of the flag tag (approximately 6 kDa) (Fig. 1B). Rapamycin is a bacterial macrolide initially used as an antifungal agent. However, due to its inhibitory effect on mTOR signaling, it is widely used as an autophagy inducer agent [24]. Activation of autophagy was confirmed using western blotting for assessment of LC3II/LC3I and p62 levels (Fig. 1C, D), according to routine protocol [21]. We further confirmed mTOR inhibition through decreased phosphorylation of S6K (ribosomal protein S6 kinase) (Fig. 1C, D). This data collectively showed activation of autophagy using rapamycin in primary neurons.

**Figure 1:**
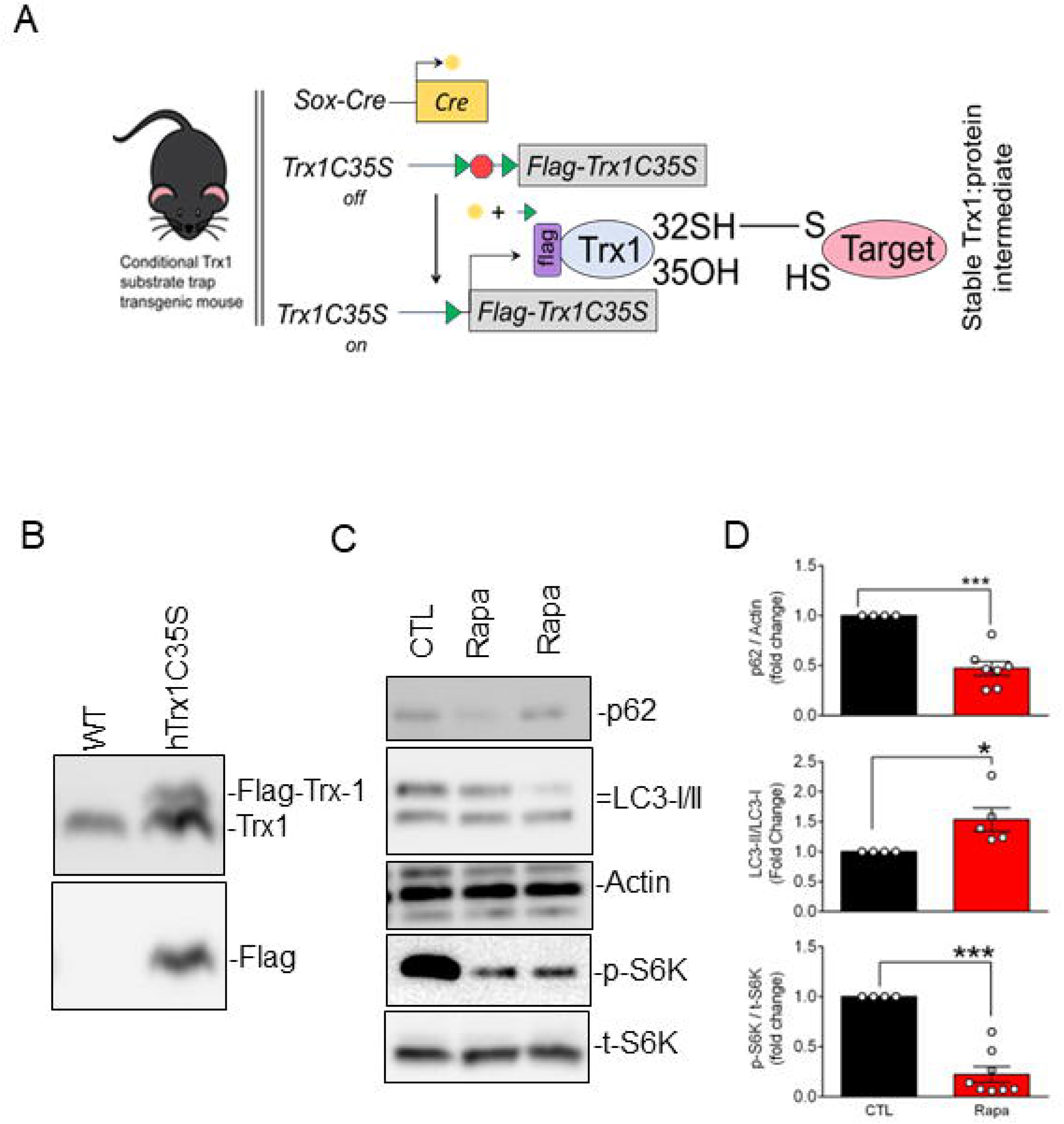
Treating Primary Neurons with Rapamycin inhibited mTOR pathway. (A) Primary cortical neurons were isolated and cultured in neurobasal-A media supplemented with N21 nutrient mixture. (B) Part of cortex were collected for genotyping for the expression of Flag-hTrx1C35S. (C) On day 8, neuronal culture was treated with 100 nM concentration of rapamycin for 24 h. Cells were then harvested for (C) immunoblot analysis of mTOR substrate after rapamycin treatment; (D) quantification of western blots shown in B. Data are mean ± SEM. Statistical significance was determined using unpaired T-test, *p<0.05 and ***p<0.001. WT – wild type; hTrx1C35S – flag tagged human Trx1C35S; CTL – control; Rapa – Rapamycin.

### Identification of neuronal Trx1 targets in mouse cortical neurons during autophagy

Rapamycin-treated neurons were harvested, and immunoprecipitation was performed to isolate the stable Trx1-targeted protein complexes using anti-flag antibody conjugated to paramagnetic beads. The immunoprecipitated samples were then washed extensively to remove the unbound proteins. The resulting co-immunoprecipitated samples were analyzed by mass spectrometry (MS) at the Manitoba Centre for Proteomics and Systems Biology as described in the materials and method section.

Using multiple biological replicates and employing stringent unique peptide cut off number, we identified 112 unique proteins (Table 1), some of which were reported previously in other systems. These include several known Trx1 substrates such as F-actin-capping protein subunit beta (CAPZB), peroxiredoxin-5 (PRDX5), cysteine and glycine rich protein 1 (CSRP1), plasminogen activator inhibitor 1 RNA-binding protein (SERBP1), as well as electrogenic sodium bicarbonate cotransporter 1 (SLC4A4) [19, 23, 25] (Table1). To investigate the role of these proteins in neuronal functions and diseases, we performed IPA (Ingenuity Pathway Analysis) and found that these proteins are involved in a wide range of neuronal functions and diseases (Table 2). A significant number of these proteins were involved in movement disorders and displayed abnormal morphology of neurons.

**Table 1:**
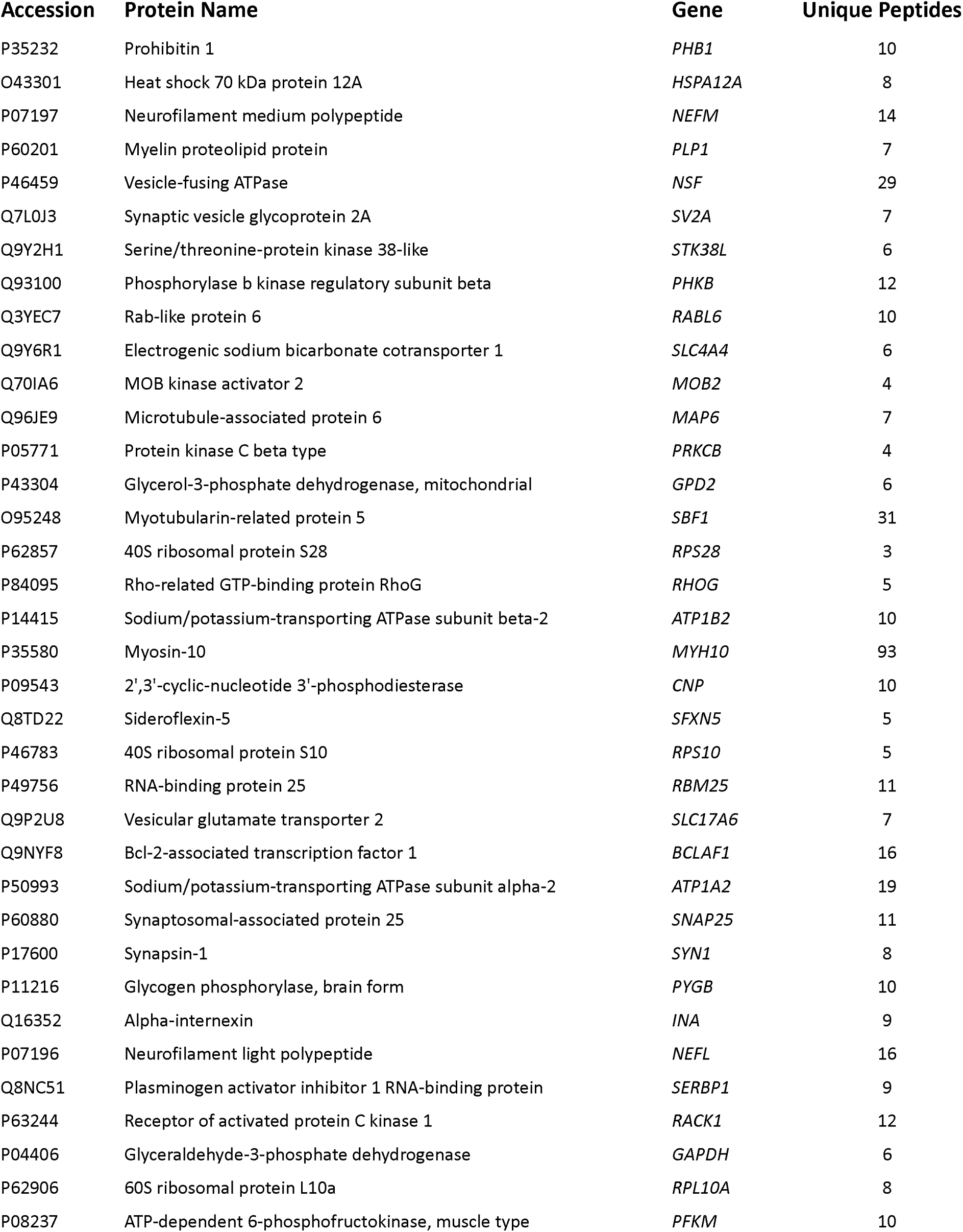
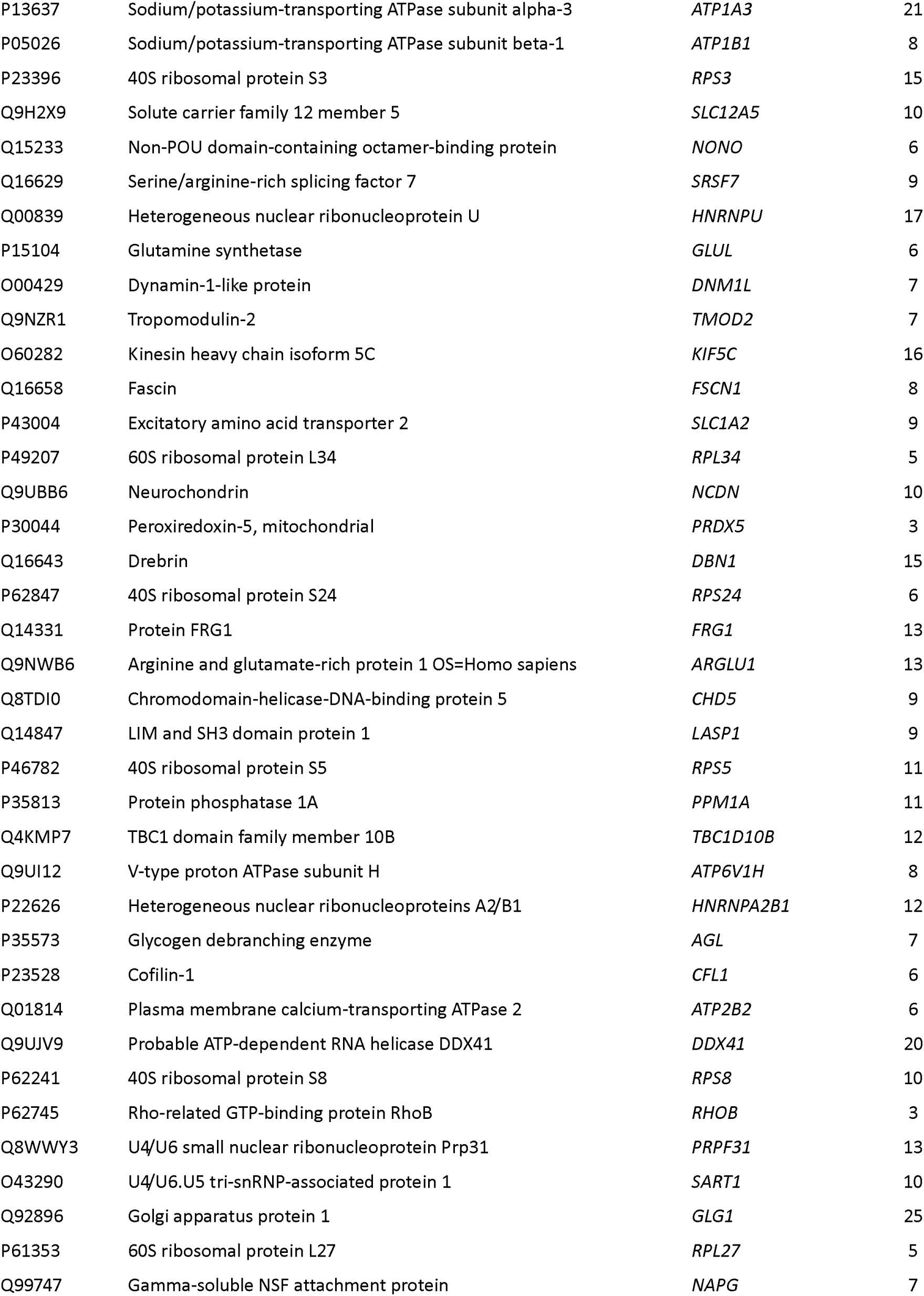
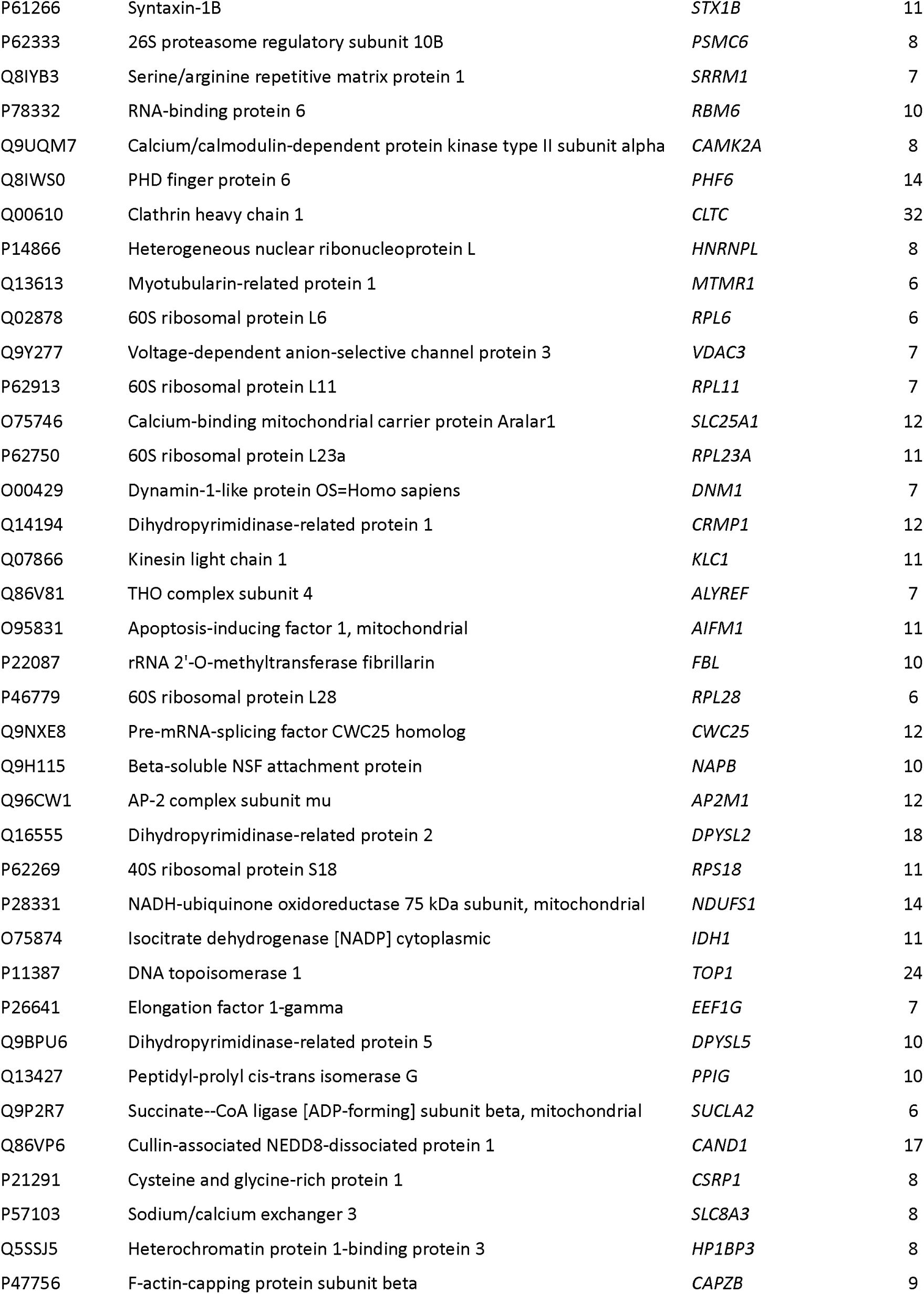
Trx1 interacting proteins in Flag-hTrx1C35S mouse primary cortical neurons.

**Table 2:**
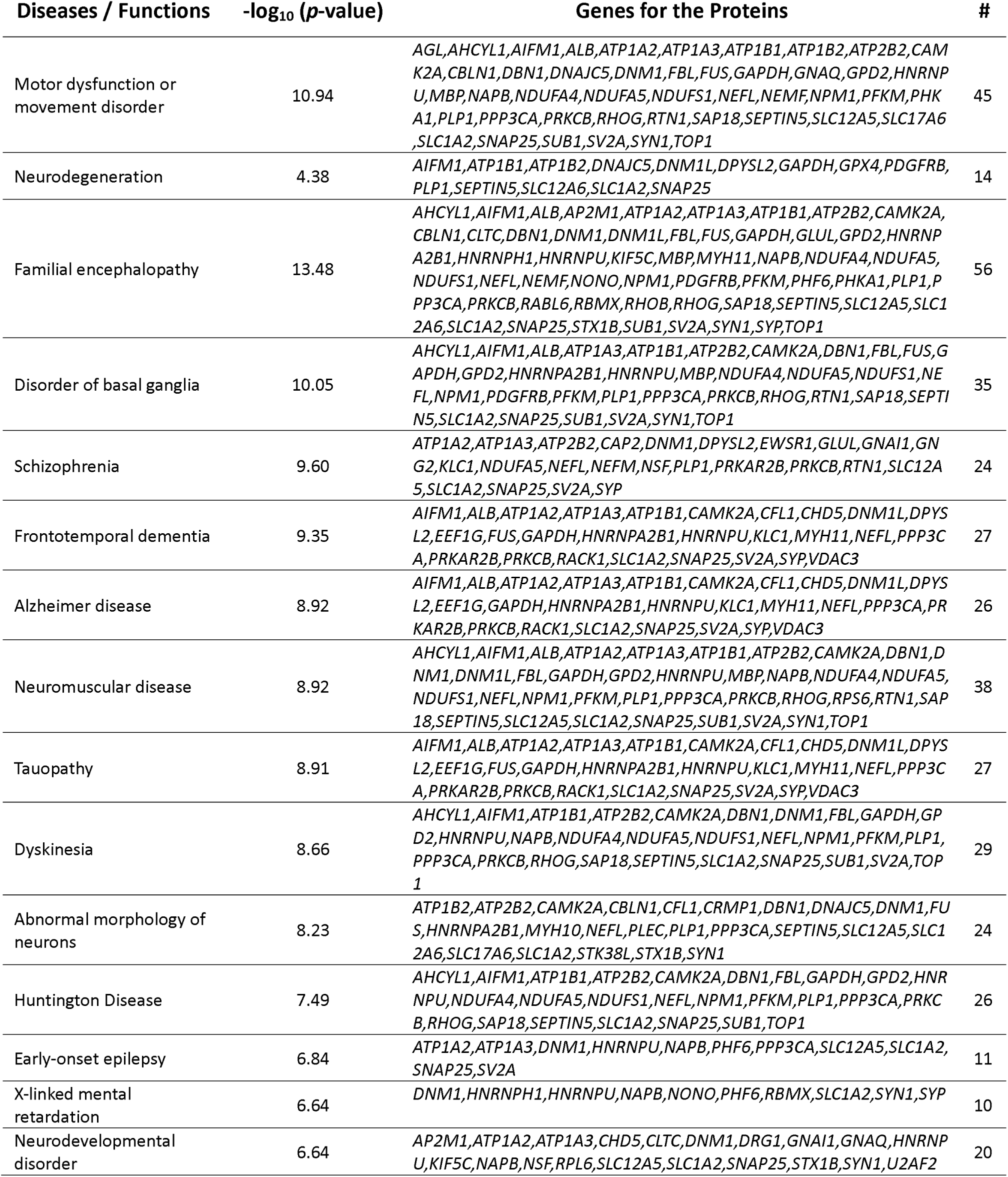
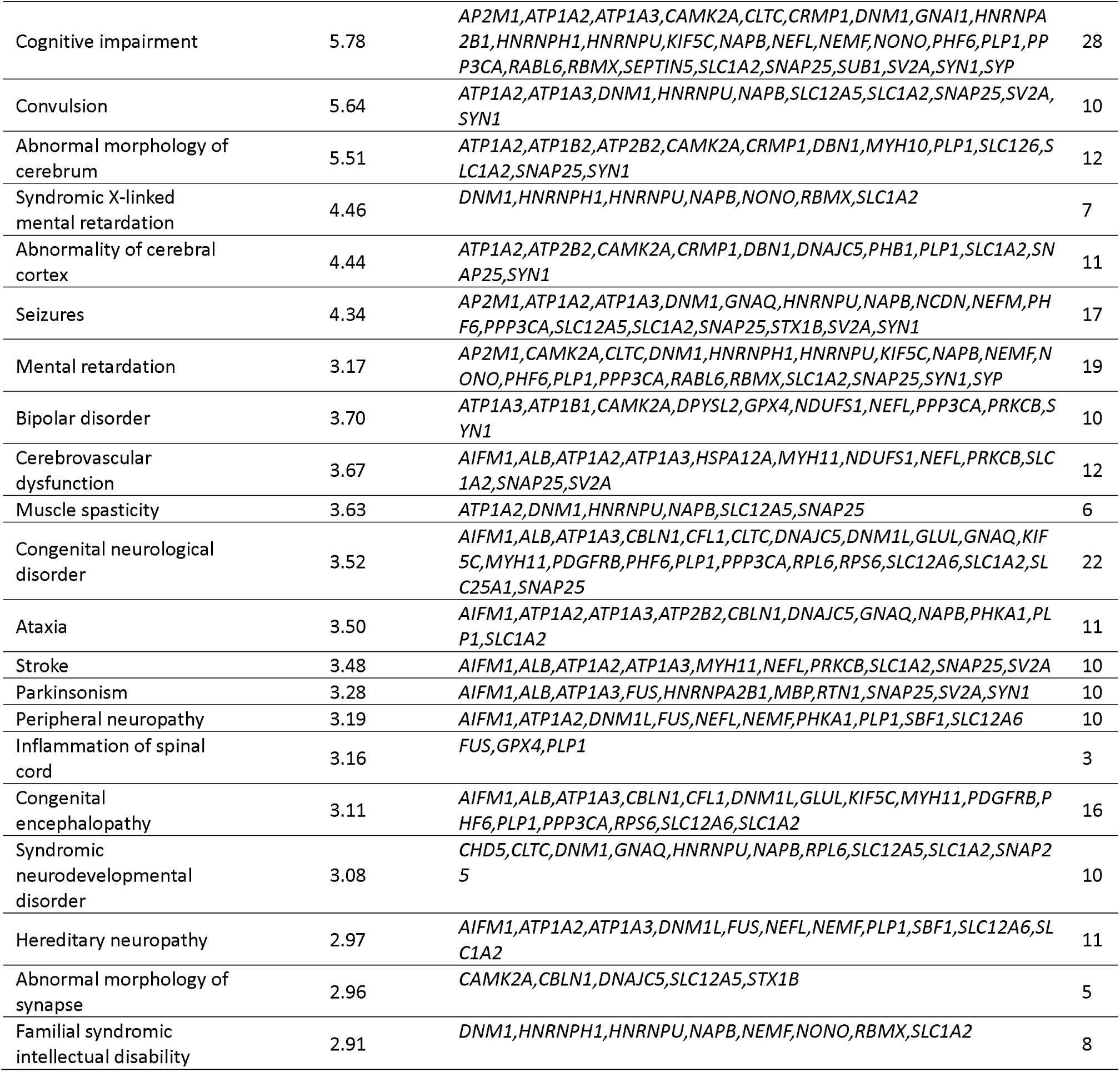
Trx1 interacting proteins involved in different neuronal diseases.

### Functional analysis of putative neuronal Trx1 targets in mouse cortical neurons

Seventy-seven Trx1-binding proteins (5 up-regulated and 72 down-regulated) were identified as significantly affected (Z score ± 2) by rapamycin treatment in mouse cortical neurons (Table 3). These proteins were analyzed using IPA to understand their roles in different cellular signaling pathways and functions. The IPA revealed the involvement of these proteins in different cellular functions and a variety of functional disorders, including ataxia and movement disorders. These proteins were also related to neuron development, neurodegeneration, and neurite outgrowth as well as those related to intracellular organellar movements including, endocytosis, microtubule dynamics, cytoskeleton organization, and cell viability (Fig. 2A). Notably, the majority of these proteins were involved in movement disorders (43), microtubule dynamics (23), and cytoskeleton organization (22) (Fig. 2B).

**Figure 2:**
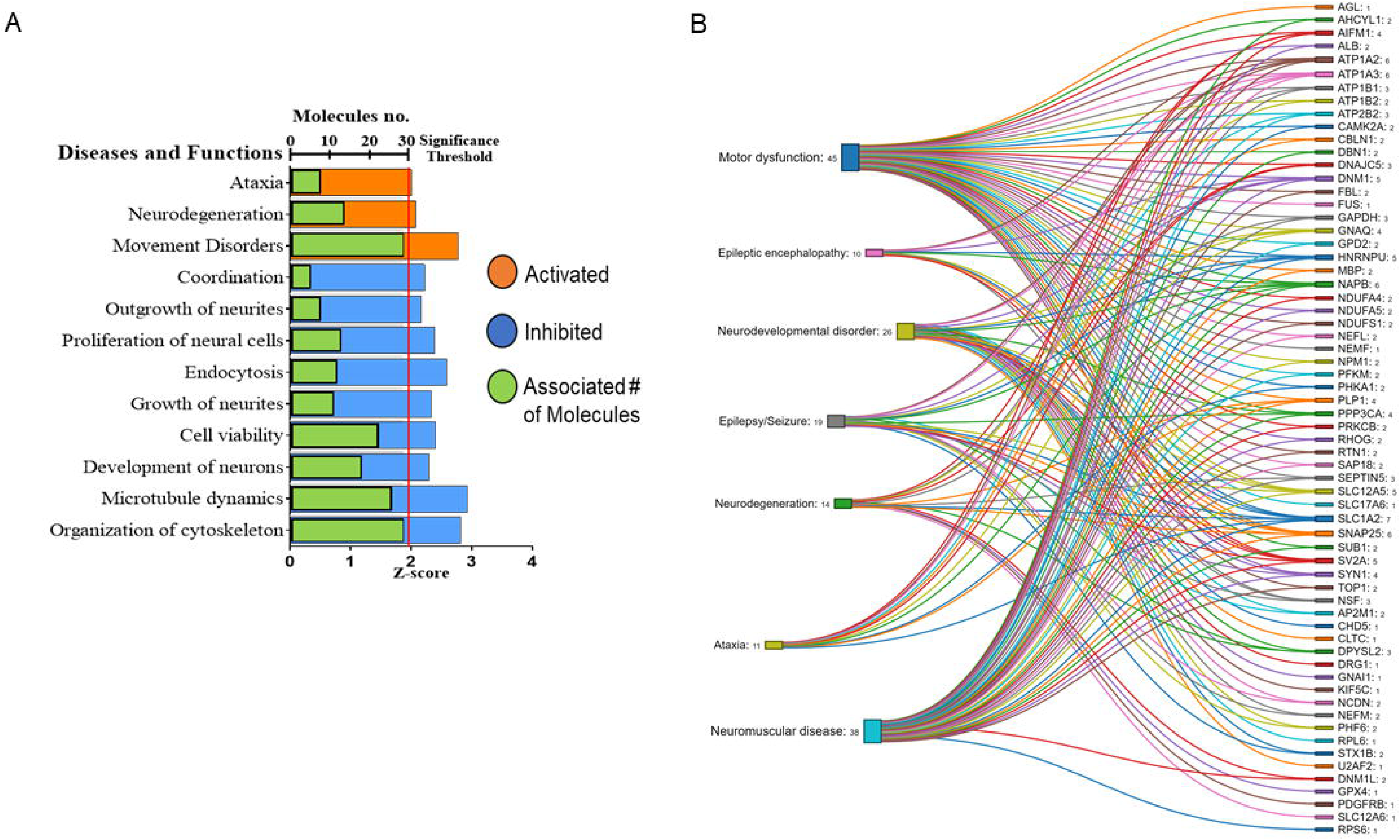
Trx1 is involved in cytoskeletal and microtubule dynamics in Rapamycin induced autophagy. (A, B) Immunoprecipitating with anti-flag antibody and proteomic analysis with Ingenuity Pathway Analysis (IPA) shows that Trx1 is involved in different cellular signaling mechanisms including cytoskeletal organization and microtubule dynamics. For (B) number right to the diseases and function denotes the proteins involved in that specific process, and number adjacent to the protein molecule indicates the number of processes the protein involved as predicted by IPA.

**Table 3:** Significantly altered proteins after rapamycin treatment.

| Accession | Protein Name | Gene | - log<sub>10</sub> (p-value) | log<sub>2</sub> Ratio |
| --- | --- | --- | --- | --- |
| P13637 | Sodium/potassium-transporting ATPase subunit alpha-3 | *ATP1A3* | 3.14 | -2.73 |
| Q16555 | Dihydropyrimidinase-related protein 2 | *DPYSL2* | 1.84 | -1.55 |
| P50993 | Sodium/potassium-transporting ATPase subunit alpha-2 | *ATP1A2* | 3.60 | -4.76 |
| P46459 | Vesicle-fusing ATPase | *NSF* | 5.50 | -3.79 |
| O95248 | Myotubularin-related protein 5 | *SBF1* | 4.36 | -2.90 |
| Q14194 | Dihydropyrimidinase-related protein 1 | *CRMP1* | 1.95 | 1.12 |
| P35813 | Protein phosphatase 1A | *PPM1A* | 2.48 | -1.33 |
| P62750 | 60S ribosomal protein L23a | *RPL23A* | 1.99 | 1.11 |
| Q07866 | Kinesin light chain 1 | *KLC1* | 1.94 | 1.08 |
| O75475 | PC4 and SFRS1-interacting protein | *PSIP1* | 1.51 | -2.33 |
| P04406 | Glyceraldehyde-3-phosphate dehydrogenase | *GAPDH* | 3.20 | -1.97 |
| P07197 | Neurofilament medium polypeptide | *NEFM* | 5.64 | -3.94 |
| Q01814 | Plasma membrane calcium-transporting ATPase 2 | *ATP2B2* | 2.37 | -2.19 |
| P43004 | Excitatory amino acid transporter 2 | *SLC1A2* | 2.72 | -3.05 |
| P11216 | Glycogen phosphorylase, brain form | *PYGB* | 3.56 | -2.94 |
| O95831 | Apoptosis-inducing factor 1, mitochondrial | *AIFM1* | 1.88 | -2.80 |
| Q9P2U8 | Vesicular glutamate transporter 2 | *SLC17A6* | 3.66 | -5.52 |
| P07196 | Neurofilament light polypeptide | *NEFL* | 3.48 | -2.08 |
| P06733 | Alpha-enolase | *ENO1* | 2.09 | -2.45 |
| P62269 | 40S ribosomal protein S18 | *RPS18* | 1.73 | -1.48 |
| P60880 | Synaptosomal-associated protein 25 | *SNAP25* | 3.58 | -2.93 |
| Q9H115 | Beta-soluble NSF attachment protein | *NAPB* | 1.86 | -2.77 |
| P17600 | Synapsin-1 | *SYN1* | 3.57 | -3.98 |
| P62847 | 40S ribosomal protein S24 | *RPS24* | 2.61 | 1.44 |
| Q93100 | Phosphorylase b kinase regulatory subunit beta | *PHKB* | 4.92 | -4.83 |
| Q05193 | Dynamin-1 | *DNM1* | 1.96 | -2.84 |
| Q9UBB6 | Neurochondrin | *NCDN* | 2.64 | -2.92 |
| P28331 | NADH-ubiquinone oxidoreductase 75 kDa subunit, mitochondrial | *NDUFS1* | 1.72 | -1.92 |
| P05771 | Protein kinase C beta type | *PRKCB* | 4.41 | -3.23 |
| Q3YEC7 | Rab-like protein 6 | *RABL6* | 4.91 | -3.34 |
| P14415 | Sodium/potassium-transporting ATPase subunit beta-2 | *ATP1B2* | 4.00 | -4.82 |
| Q4KMP7 | TBC1 domain family member 10B | *TBC1D10B* | 2.46 | -3.27 |
| O75367 | Core histone macro-H2A.1 | *MACROH2A1* | 1.78 | -2.74 |
| Q14847 | LIM and SH3 domain protein 1 | *LASP1* | 2.53 | -1.80 |
| P63096 | Guanine nucleotide-binding protein G(i) subunit alpha-1 | *GNAI1* | 2.50 | -1.37 |
| Q16352 | Alpha-internexin | *INA* | 3.53 | -3.79 |
| Q96CW1 | AP-2 complex subunit mu | *AP2M1* | 1.85 | -1.11 |
| P05026 | Sodium/potassium-transporting ATPase subunit beta-1 | *ATP1B1* | 3.12 | -2.83 |
| Q9Y6R1 | Electrogenic sodium bicarbonate cotransporter 1 | *SLC4A4* | 4.55 | -4.36 |
| O43301 | Heat shock 70 kDa protein 12A | *HSPA12A* | 5.97 | -4.02 |

| Accession | Protein Name | Gene | - log<sub>10</sub> (*p*-value) | log<sub>2</sub> Ratio |
| --- | --- | --- | --- | --- |
| Q9Y2H1 | Serine/threonine-protein kinase 38-like | *STK38L* | 5.08 | -4.30 |
| Q96JE9 | Microtubule-associated protein 6 | *MAP6* | 4.52 | -3.67 |
| Q13613 | Myotubularin-related protein 1 | *MTMR1* | 2.04 | -2.57 |
| Q99747 | Gamma-soluble NSF attachment protein | *NAPG* | 2.26 | -2.60 |
| P84095 | Rho-related GTP-binding protein RhoG | *RHOG* | 4.07 | -4.54 |
| Q7L0J3 | Synaptic vesicle glycoprotein 2A | *SV2A* | 5.16 | -3.96 |
| O00429 | Dynamin-1-like protein | *DNM1L* | 2.81 | -2.46 |
| P43304 | Glycerol-3-phosphate dehydrogenase, mitochondrial | *GPD2* | 4.40 | -3.34 |
| Q08209 | Protein phosphatase 3 catalytic subunit alpha | *PPP3CA* | 2.18 | -2.05 |
| P46779 | 60S ribosomal protein L28 | *RPL28* | 1.87 | 0.97 |
| Q92930 | Ras-related protein Rab-8B | *RAB8B* | 3.43 | -2.34 |
| Q9Y295 | Developmentally-regulated GTP-binding protein 1 | *DRG1* | 2.63 | -3.24 |
| P62745 | Rho-related GTP-binding protein RhoB | *RHOB* | 2.30 | -1.90 |
| Q99719 | Septin-5 | *SEPTIN5* | 3.09 | -2.99 |
| Q9H3Z4 | DnaJ homolog subfamily C member 5 | *DNAJC5* | 2.30 | -1.34 |
| P63010 | AP-2 complex subunit beta | *AP2B1* | 1.69 | -2.04 |
| Q16799 | Reticulon-1 | *RTN1* | 3.11 | -2.10 |
| Q70IA6 | MOB kinase activator 2 | *MOB2* | 4.52 | -4.18 |
| P11498 | Pyruvate carboxylase, mitochondrial | *PC* | 1.63 | -1.86 |
| O60524 | Ribosome quality control complex subunit NEMF | *NEMF* | 2.22 | -2.45 |
| P30044 | Peroxiredoxin-5, mitochondrial | *PRDX5* | 2.63 | -1.29 |
| O43759 | Synaptogyrin-1 | *SYNGR1* | 1.68 | -2.98 |
| O00422 | Histone deacetylase complex subunit SAP18 | *SAP18* | 1.71 | -1.68 |
| P36969 | Phospholipid hydroperoxide glutathione peroxidase | *GPX4* | 2.99 | -2.80 |
| O00483 | Cytochrome c oxidase subunit NDUFA4 | *NDUFA4* | 1.88 | -1.89 |
| Q9UHW9 | Solute carrier family 12 member 6 | *SLC12A6* | 1.76 | -2.53 |
| P08195 | 4F2 cell-surface antigen heavy chain | *SLC3A2* | 2.60 | -1.94 |
| P37108 | Signal recognition particle 14 kDa protein | *SRP14* | 1.98 | -1.76 |
| Q96CT7 | Coiled-coil domain-containing protein 124 | *CCDC124* | 2.51 | -4.06 |
| Q5JTH9 | RRP12-like protein | *RRP12* | 2.89 | -1.21 |
| Q9H598 | Vesicular inhibitory amino acid transporter | *SLC32A1* | 1.54 | -3.02 |
| Q16718 | NADH dehydrogenase [ubiquinone] 1 alpha subcomplex subunit 5 | *NDUFA5* | 2.13 | -1.41 |
| P09619 | Platelet-derived growth factor receptor beta | *PDGFRB* | 1.81 | -0.90 |
| Q9NQ50 | 39S ribosomal protein L40, mitochondrial | *MRPL40* | 1.46 | -3.63 |
| Q8NBX0 | Saccharopine dehydrogenase-like oxidoreductase | *SCCPDH* | 3.01 | -1.59 |
| P31944 | Caspase-14 | *CASP14* | 2.75 | -2.42 |
| P22234 | Bifunctional phosphoribosylaminoimidazole carboxylase | *PAICS* | 1.45 | -2.77 |

Changes in cellular morphology and cytoplasmic transport of cellular organelles are commonly observed in autophagy. Accordingly, IPA showed that Trx1-interacting proteins were involved in formation, organization, and stabilization of the cytoskeleton by modulating microtubule dynamics, through polymerization and depolymerization of actin filaments (Fig. 3A). Additionally, IPA predicted that rapamycin treatment impacts various signaling pathways in mouse cortical neurons; most significantly those related to neuronal function include mTOR signaling, mitochondrial dysfunction, regulation of lysosomal membranes, Rho GTPase signaling, Glycolysis I, AMPK signaling, etc. (Table 4). We further focused on the protein network in mTOR signaling and the Rho GTPase family (Fig. 3B and Fig. 4) which showed that Trx1 interacting proteins are involved in actin organization by affecting actin polymerization and nucleation.

**Figure 3:**
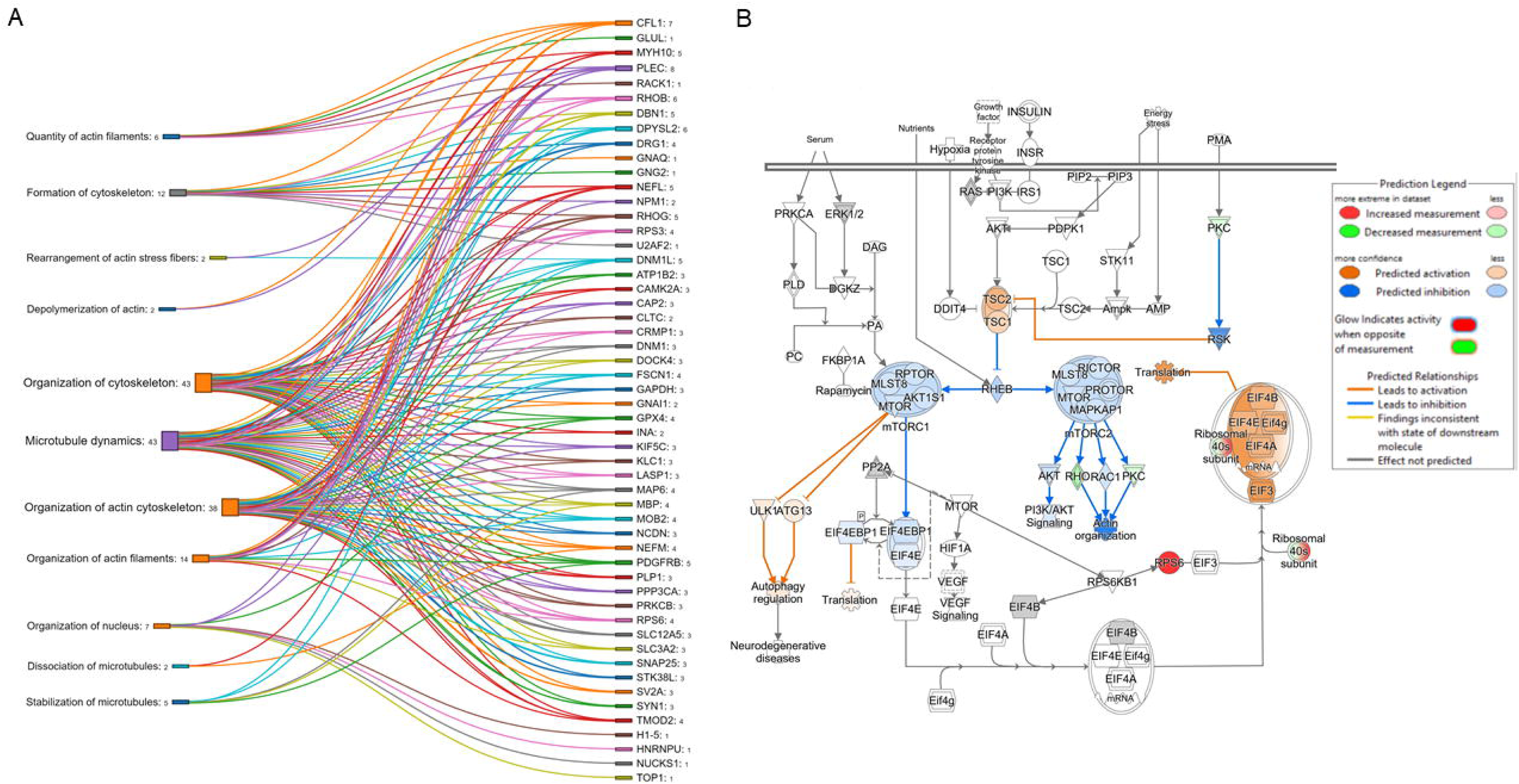
Impact of Trx1 interacting protein after Rapamycin treatment on cytoskeleton dynamics. (A) Number of Trx1 interacting proteins identified by IPA in different cytoskeleton organizations. Number adjacent to the biological processes are the number of proteins associated with the specific process, number adjacent to the protein denotes the number of processes that protein involved in. (B) Association of proteins predicted by IPA in cellular signaling pathways affecting autophagy, translation, and actin organization.

**Figure 4:**
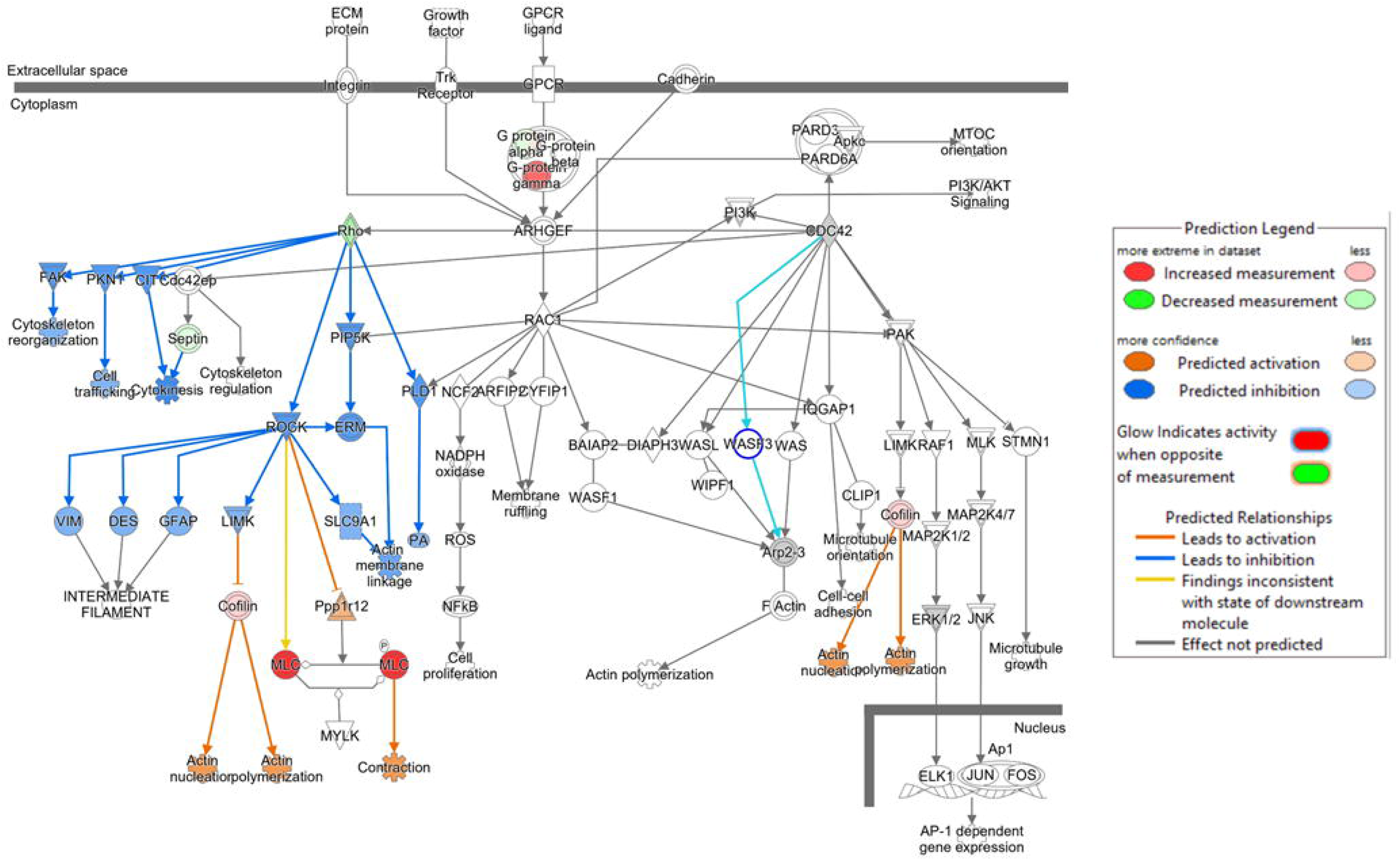
Rho Signaling pathways and actin dynamics predicted by IPA analysis of Trx1 interacting pathways. The blue line predicted potential inhibition and brown line predicted possible activation of the signaling pathway as annotated by IPA. The colored nodes represent Trx1 interacting proteins and unshaded nodes indicate additional proteins suggested by IPA.

**Table 4:** Proteins involved in various signaling pathways predicted by IPA.

| Category | $-\log_{10}(p\text{-value})$ | Genes for the Proteins Involved |
| --- | --- | --- |
| EIF2 Signaling | 15.8 | *RPL10A, RPL11, RPL18, RPL19, RPL22L1, RPL23A, RPL27, RPL28, RPL34, RPL35A, RPL37A, RPL6, RPS10, RPS18, RPS20, RPS24, RPS28, RPS3, RPS6, RPS8* |
| mTOR Signaling | 6.48 | *PRKCB, RHO B, RHOG, RPS10, RPS18, RPS20, RPS24, RPS28, RPS3, RPS6, RPS8* |
| Tight Junction Signaling | 5.32 | *MYH10, MYH11, MYL6, NAPB, NAPG, NSF, PRKAR2B, SNAP25, STX1B* |
| Synaptogenesis Signaling Pathway | 4.86 | *AP2M1, CAMK2A, CFL1, DNAJC5, NAPB, NAPG, NSF, PRKAR2B, SNAP25, STX1B, SYN1* |
| Mitochondrial Dysfunction | 4.56 | *AIFM1, GPD2, GPX4, NDUFA4, NDUFA5, NDUFS1, PRDX5, VDAC3* |
| Protein Kinase A Signaling | 4.48 | *CAMK2A, GNAI1, GNAQ, GNG2, H15, MYH10, MYL6, PHKB, PPP3CA, PRKAR2B, PRKCB, PYGB* |
| Regulation of eIF4 and p70S6K Signaling | 4.38 | *RPS10, RPS18, RPS20, RPS24, RPS28, RPS3, RPS6, RPS8* |
| Signaling by Rho Family GTPases | 3.95 | *CFL1, GNAI1, GNAQ, GNAZ, GNG2, MYL6, RHO B, RHOG, SEPTIN5* |
| Phagosome Maturation | 3.91 | *ATP6V1H, NAPB, NAPG, NSF, PRDX5, SNAP25, STX1B* |
| Sphingosine-1-phosphate Signaling | 3.7 | *CASP14, GNAI1, GNAQ, PDGFRB, RHO B, RHOG* |
| nNOS Signaling in Neurons | 3.49 | *CAMK2A, PFKM, PPP3CA, PRKCB* |
| Glycolysis I | 3.06 | *ENO1, GAPDH, PFKM* |
| Glutamate Receptor Signaling | 2.93 | *GLUL, GNG2, SLC17A6, SLC1A2* |
| AMPK Signaling | 2.8 | *GNAI1, GNAQ, GNAZ, GNG2, PFKM, PPM1A, PRKAR2B* |
| Gap Junction Signaling | 2.57 | *DBN1, GNAI1, GNAQ, PPP3CA, PRKAR2B, PRKCB* |
| Glycogen Degradation II | 2.52 | *AGL, PYGB* |
| HIF1α Signaling | 1.81 | *CAMK2A, PPP3CA, PRKCB, RACK1, RPS6* |

### Validation of approach for identifying neuronal Trx1 targets

Western blot data revealed a significant inhibition of S6 phosphorylation in primary cortical neurons treated with rapamycin, indicating a blockade of the mTOR signaling pathway (Fig. 5A). Proteomic analysis suggested an overall decrease for RhoB and RhoG in rapamycin treated group compared to the control group (Table 3). Inhibition of mTOR by rapamycin treatment increased RhoB expression levels and decreased RhoG level compared to the control group (Fig. 5A). Although the RhoB protein level was increased in total cell lysate after rapamycin treatment, the flag antibody pulled RhoB was found to be decreased in mass spectrometry analysis (Table 3). This difference in RhoB could be due to altered binding and interaction of Trx1 in rapamycin treated samples which required further investigation. Interaction of Trx1 with RhoB in normal conditions was confirmed using immunoprecipitation (Fig. 5B). Previous studies have shown that mTOR increases globular actin (G-actin) polymerization and/or decreases filamentous actin (F-actin) depolymerization [26], and rapamycin inhibits actin fibrilization. Hence, we analyzed the actin polymerization status in rapamycin-treated neurons by measuring the ratio of F/G actin. As per the previous reports conducted in other cell types, induction of autophagy in neurons decreased F/G actin ratio indicating the reduction of actin polymerization (Fig. 5A). To further substantiate our findings, we labelled primary cortical neurons with actin-phalloidin antibody and compared rapamycin treated and untreated neurons. Confocal microscopy data revealed that rapamycin treatment reduced the fibrillar actin puncta across cortical neuronal axons compared to untreated cortical neurons (Fig. 5C).

**Figure 5:**
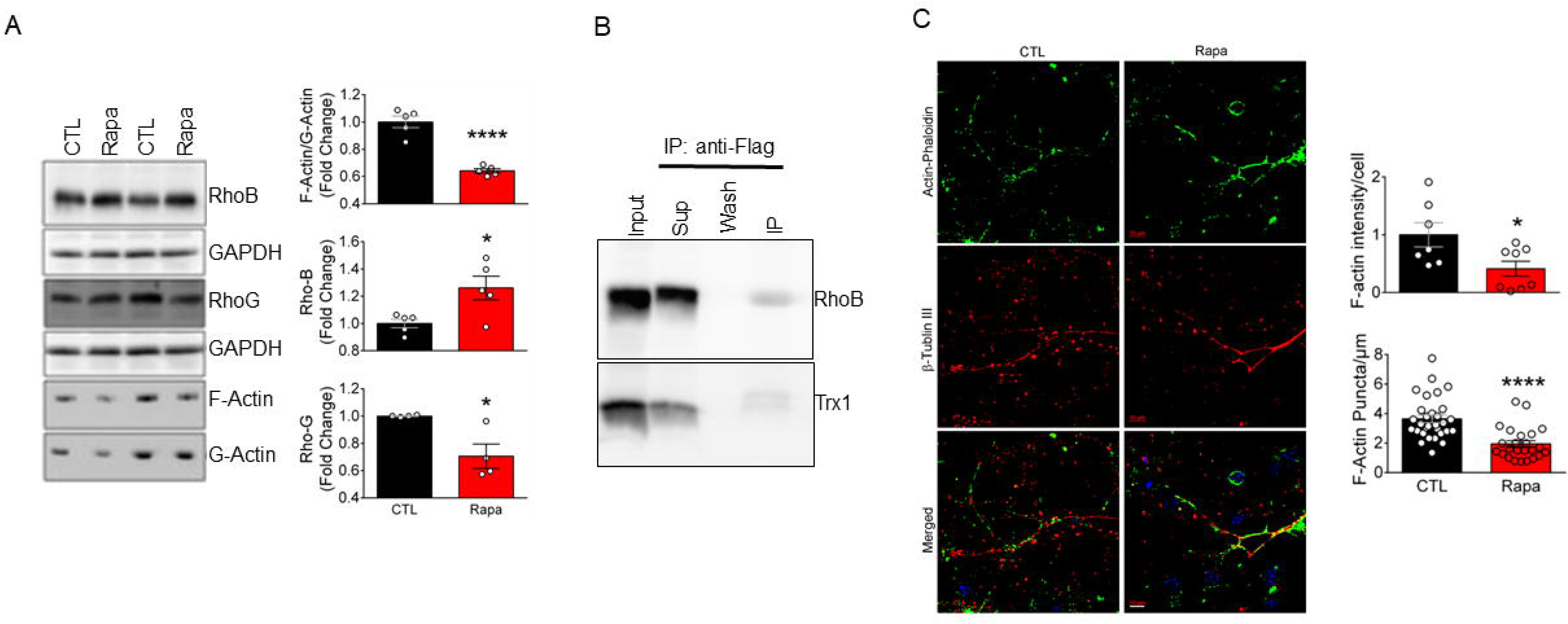
Inhibition of mTOR was associated with decreased actin polymerization. (A) Western blot analysis showed that rapamycin increased RhoB expression, decreased RhoG expression and reduced the fibrillar actin in primary cortical neurons. Bar graph showing densitometry analysis of western blot bands, each circle represents a biological replicate. (B) Immunoprecipitation with anti-Flag M2 antibody detected both RhoB and Trx1 in primary cortical neurons. Input = cell lysate, sup = supernatant collected after interaction with flag bead. (C) Confocal microscopy revealed that rapamycin treatment reduced the actin filament across cortical neuronal axons compared to untreated cortical neurons. Primary cortical neurons were immunolabeled with actin-phaloidin and β-tubulin III. Scale bar = 10 µm. Data are the mean ± SEM, * p<0.05, **** p<0.001. CTL – control; Rapa – Rapamycin.

### The key role of Trx1 in actin fibrilization in autophagy

Proteomic data indicated a direct interaction between Trx1 and RhoB and RhoG, we aimed to further evaluate the importance of Trx1 by generating a Trx1 knockdown (Trx1KD) in HT22 hippocampal cells. To induce autophagy, we exposed the scrambled control (SCM) and Trx1KD-HT22 cells to serum deprivation (SD) for 24 hours. Activation of autophagy and actin reorganization were tested by phosphorylation of S6-ribosomal protein and the F/G actin ratio, respectively. Inhibition of S6-ribosomal protein phosphorylation was successfully achieved in both SCM and Trx1KD cells (Fig. 6A). SD also decreased F/G actin ratio in control SCM cells, however, Trx1KD controls had significantly lower F/G actin ratio than the control SCM.

**Figure 6:**
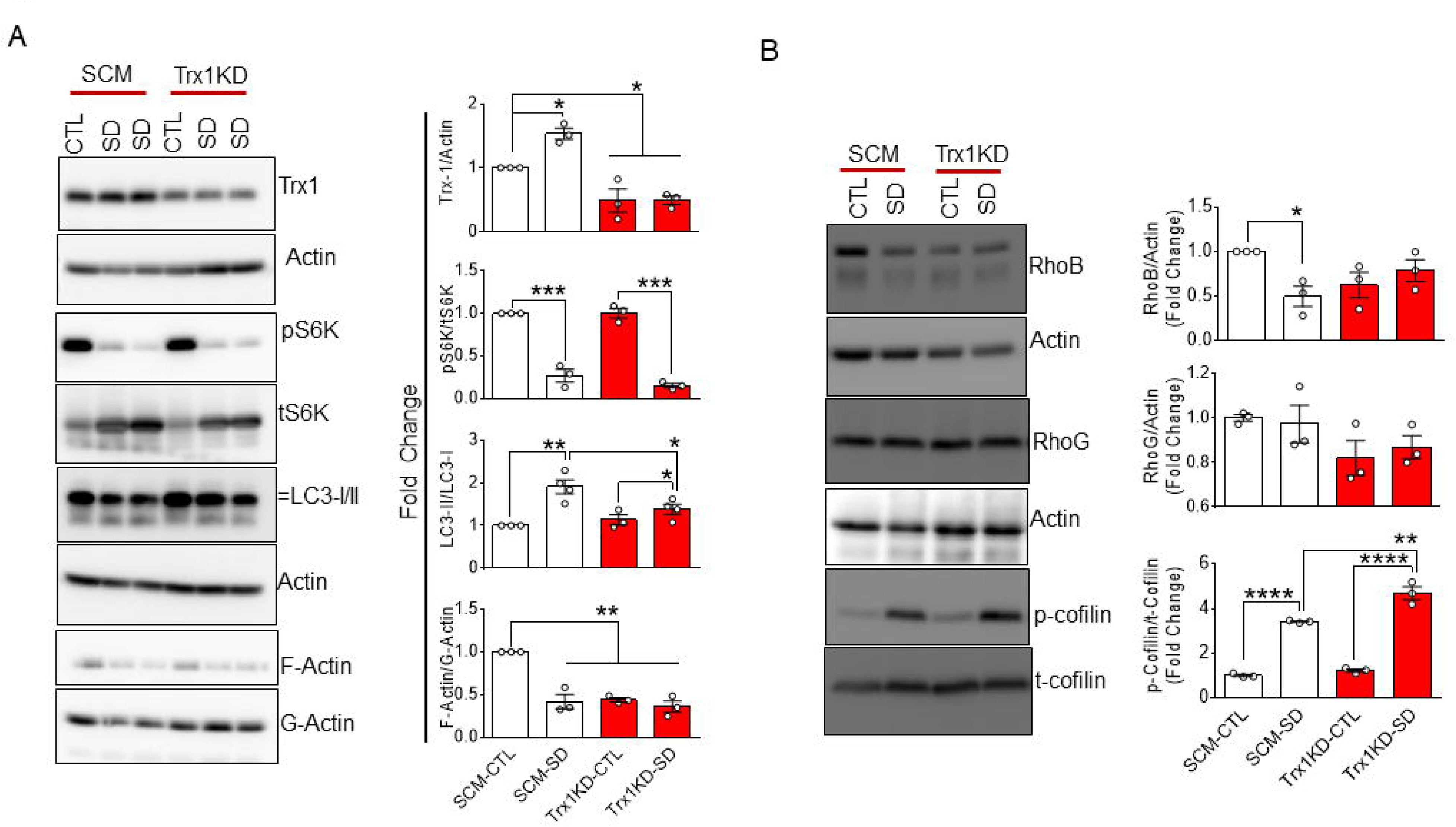
Knocked down of Trx1 in HT22 cells resulted in decreased actin polymerization. (A) Scrambled control (SCM) and Trx1 Knocked Down (Trx1KD HT22) cells were subjected to serum deprivation for 24 h. Cell lysate was subjected to western blot for the indicated markers. Densitometry analysis showed inhibition of mTOR pathway after serum deprivation. Only Trx1 knock down was sufficient to reduce the F-actin in HT22 cells. Each circle indicates a biological replicate. (B). RhoB and RhoG did not significantly change in Trx1KD HT22 cells. Phosphorylation of cofilin was only observed in serum deprived conditions regardless of Trx1KD. Data are presented as mean ± SEM. * p<0.05, ** p<0.01, **** p<0.0001. CTL – control; SD – serum deprivation.

Additionally, SD in Trx1KD cells did not exacerbate the F/G Actin ratio (Fig. 6A), indicating the pivotal role of Trx1 in actin polymerization in normal physiological conditions (Fig. 6A). To find out whether actin depolymerization is Rho GTPase dependent, immunoblotting was performed for RhoB and RhoG. No significant changes in RhoB and RhoG expression were observed across the experimental groups in Trx1KD-HT22 cells in control condition (Fig. 6B). To further investigate if the reduced levels of F-Actin in Trx1KD cells was cofilin dependent, we examined the phosphorylation of cofilin in SCM and Trx1KD-HT22 cells. The cofilin phosphorylation levels were comparable in both SCM and TrxKD-HT22 cells in normal conditions and significantly increased when SCM and Trx1KD HT22 cells were exposed to SD (Fig. 6B).

The impact of Trx1 on actin fibrilization was further assessed by the confocal microscopy after labeling the cells with actin-phalloidin. The images were analyzed by the ridge detection plugin in ImageJ to determine the average length of the filaments per cell area and the average number of filaments per cell area. A minimum of 30 cells were randomly selected from each group in three different biological replicates. We observed a reduction of length of filaments per cell area in Trx1KD HT22 control cells when compared to SCM, while Trx1KD cells had increased number of small filaments in comparison to SCM (Fig. 7A). The role of Trx1 in actin fibrilization was also tested in primary cultures of rat cortical neurons (Fig. 7B).

**Figure 7:**
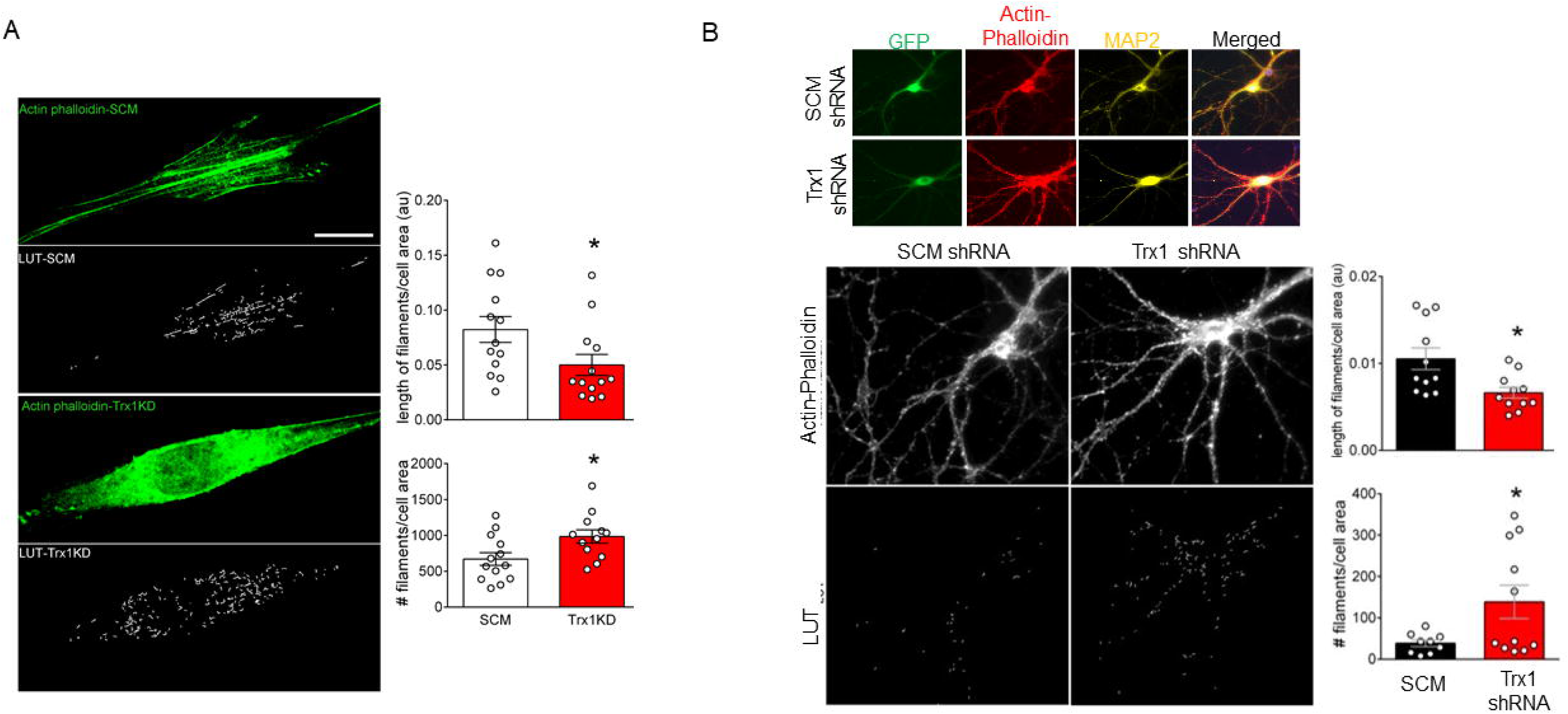
Actin polymerization and fibrilization is reduced in Trx1KD cells. (A) SCM and Trx1KD HT22 cells were immune-stained with actin-phaloidin to visualize the F-actin through confocal microscope. Length of actin filaments and number of actin filaments per cell area was measured using Ridge Detection Plugin in ImageJ. Data indicates mean ± SEM, * p<0.05. (B) Rat primary cortical neurons were transfected with either LV-shRNA-scramble or LV-shRNA-Trx1 through nucleofection. Fixation and immunostaining of neurons were carried out on DIV 18. F-actin was assayed by immunostaining of actin phalloidin co-labeled with MAP2 in scramble or Trx1 shRNA treated cortical neurons using Ridge Detection Plugin in ImageJ. Data are mean ± SEM, ** p<0.01. SCM – scrambled control, Trx1KD – Trx1 knocked down; LUT – look up table; # - number.

## Discussion

Through the study of putative neuronal targets of Trx1 in mouse cortical neurons, we have gained valuable insight into the complex regulatory roles of Trx1 that govern cellular signaling pathways including cytoskeletal organization and actin dynamics. Our findings highlight the multifunctional roles of Trx1 in maintaining cellular homeostasis, many of which remain unknown and are mediated by the simple redox regulatory role of Trx1. We propose that Trx1 orchestrates the cellular proteins’ performance like a “behind the curtain” conductor.

By utilizing the flag-tagged hTrx1C35S mutant, stable mixed disulfide adducts were obtained, thus enabling proteomic analysis for identifying Trx1 interacting proteins. The discovery of 112 distinct proteins, out of which 77 displayed significant changes after rapamycin treatment, highlights the complex nature of Trx1 involvement in cellular signaling pathways. Previous reports in these animals have showed that expression of flag-hTrx1C35S does not alter the expression of endogenous thioredoxin and glutathione systems [23]. Also, hTrx1C35S does not show catalytic activity, nor does it affect the redox activity of endogenously expressed Trx1 [23]. Therefore, we believe that this model reflects the extent of Trx1 involvement in redox regulation of these proteins.

The potential targets of Trx1 were analyzed for their function using IPA software, indicating their involvement in various cellular processes such as mTOR signaling, mitochondrial dysfunction, lysosomal regulation, and Rho GTPase signaling. These findings are consistent with previous reports on the diverse roles of Trx1[19, 25, 27]. The study also found that Trx1 plays a critical role in regulating cytoskeletal organization in normal cellular conditions which is independent of the Rho signaling pathway. Rho family proteins are known to regulate the actin organizations by controlling actin polymerization and depolymerization through cofilin phosphorylation [28, 29]. Enhanced G-actin level was detected in Trx1KD cells even under normal conditions, we did not observe any significant changes in Rho proteins and cofilin phosphorylation. This is in line with previous reports showing the interaction of actin with Trx1 and eventually in inhibition of F-actin formation [30, 31]. These studies have shown that Trx1 interacts with G-actin under oxidative stress through Cys62 and prevents the formation of F-actin. However, in this study we have observed that knocking down Trx1 in HT22 cells also increased the G-actin compared to F-actin. This may indicate that in physiological condition Trx1 plays a role in actin organization by mediating signaling pathways responsible for actin fibrilization and depolymerization. Further studies are required to determine the direct involvement of Trx1 in actin dynamics. Our mass spectroscopy data also showed several proteins which play a crucial role in actin cytoskeletal maintenance such as CAPZB, which is responsible for barbing the actin filament [32]. Alpha-internexin (INA) is a type of Class-IV neuronal intermediate filament that can self-assemble and plays a crucial role in the morphogenesis of neurons. INA may form an independent structural network, or it may work together with NF-L (Nerofilament – light) to create the filamentous backbone. NF-M and NF-H (Neurofilament - medium and heavy) can then attach to the backbone to form cross-bridges [33]. Synapsin-1 (SYN1) is a Trx1-binding phosphoprotein that is predominantly found in neurons. SYN1 primary function is to coat synaptic vesicles and bind to the cytoskeleton. It has been reported that vesicle release is controlled by membrane potential [34], our proteomic data showed several proteins which are responsible for maintaining membrane potential, thereby could play crucial roles in actin organization. For example, Solute carrier family 12 member 5 (SLC12A5) mediates the movement of potassium and chloride ions across cell membranes in mature neurons, which is essential for maintaining proper levels of chloride ions within neurons. Removing excess chloride from inside neurons ensures that low levels of chloride required for proper function are maintained [35]. We also identified excitatory amino acid transporter 2 (SLC1A2), vesicular glutamate transporter 2 (SLC17A6), electrogenic sodium bicarbonate cotransporter 1 (SLC4A4), solute carrier family 12 member 6 (SLC12A6), vesicular inhibitory amino acid transporter (SLC32A1) and few more transporters which are responsible for maintaining membrane potential. The interaction of Trx1 with these proteins warrants further investigation into the intricate regulatory network that involves Trx1, considering the unique cellular environments and possible implications for maintaining actin homeostasis. To verify the reliability of our proteomic results, we conducted Western blot analysis that confirmed the inhibition of mTOR signaling induced by rapamycin. This was evidenced by a decrease in S6 phosphorylation. This finding provided further evidence of the interplay between Trx1, mTOR, and actin polymerization and is in line with previous studies that have demonstrated the direct influence of Trx1 on mTOR activity [27].

This study unveiled a previously unrecognized role of Trx1 as a crucial catalyst in actin polymerization. Previous reports predominantly focused on various other signaling molecules, shedding light on the intricate network of intracellular interactions necessary for actin-fibril formation. The identification of Trx1’s involvement enriches our comprehension of the molecular underpinnings governing cytoskeletal dynamics and opens new avenues for exploring the regulation of cellular architecture and motility. Given the essential functions of actin structures in numerous cellular processes, from movement to division, our findings suggest that Trx1 could play a critical role in a wide array of physiological and pathological contexts. The detailed mechanism through which Trx1 influences actin polymerization remains an exciting frontier for further investigation, promising to unravel new layers of complexity in cell biology. The validation of our findings in primary neurons reinforces the crucial role of Trx1 in maintaining cytoskeletal stability. Our results are consistent with prior studies highlighting Trx1’s impact on actin dynamics [36, 37], laying the groundwork for comprehending the broader implications of Trx1 dysregulation in neurons.

## Conclusion

In conclusion, our study provides novel perspectives on the role of Trx1 in neuronal signaling pathways, mTOR activation, and actin dynamics. By pinpointing Trx1 targets and verifying them in different cellular models, this study advances our understanding of the diverse roles Trx1 in maintaining neuronal homeostasis. Further research is needed to investigate the role of Trx1 in actin dynamics in the nervous system.

## Experimental procedures

### Materials

The following materials are used for this study: Dulbecco’s Modified Eagle Medium (DMEM) High Glucose, Opti-MEM (Minimum Essential Media) Reduced Serum Media, Trypsin, GlutaMAX supplement (100X), and Penicillin-Streptomycin-Neomycin (PSN) Antibiotic Mixture (10,000 U/mL), Dulbecco′s Phosphate Buffered Saline (DPBS), Fetal bovine serum (FBS) was purchased from GibcoTM (Life Technologies, Canada). Lipofectamine RNAiMax transfection reagent was from Invitrogen, USA. Scramble (LV-shRNA-scramble, #LVP015-G) and Trx1 shRNA plasmids (LV-shRNA-Trx1, #488660940395) were purchased from Applied Biological Materials Inc., Canada. Pierce™ BCA Protein Assay Kit and Halt™ Protease and Phosphatase Inhibitor Single-Use Cocktail were from Thermo Scientific™, USA. The Western blotting detection kit was from Bio-rad Laboratories Inc., USA. Glass cover slips were from Fisher Scientific, Canada.

### Primary Neuronal Culture

Neurons were cultured from flag-hTrx1C35S mice embryos on E17 according to routine procedures [38]. Cerebral Cortex from each embryo was dissected and kept in artificial cerebrospinal fluid (aCSF) on ice. The tissue from each embryo was separately cultured as follows: the cortex was washed twice with aCSF, resuspended in aCSF, containing papain (1:1000), and incubated at 37°C for 15 min with intermittent gentle shaking. To neutralize the protease activity, an equal volume of FBS was added to aCSF, and the mixture was centrifuged at 900 rpm for 5 min. The supernatant was discarded, and the pellets were resuspended in aCSF with 1X DNAase (Sigma# D5025). The cells were triturated with a polished Pasteur pipette 20 times and centrifuged at 900 rpm for 5 min. The pelleted cells were resuspended in Neurobasal A media, containing 1% N21 max supplement, PSN, and L-glutamine. After counting, the cells were seeded in a PDL coated 6 well plate at a density of 150,000 cells/well. Half of the growth medium was changed every 6 days. PDL-coated glass coverslips were used to seed cells for immunocytochemistry. The genotype for each individual embryo was tested using PCR and western blotting in a piece of CNS tissue to confirm the presence of Trx1 and flag proteins.

### Immunoprecipitation and Mass Spectroscopy

After confirming the genotyping, primary cortical neurons were maintained for 8 days in neurobasal-A media supplemented with N21 nutrient mixture. On day 8, neuronal cultures were treated with 100 nM concentration of rapamycin for 24 h. Cells were then harvested and immunoprecipitated with anti-flag antibody (Sigma Aldrich, M8823). For mass spectrometry (MS) proteomic analysis, the resulting co-immunoprecipitated samples were analyzed to detect interaction partners. This was completed using label-free LC-MS/MS analysis at the Manitoba Centre for Proteomics and Systems Biology. On-bead precipitates from 200 mg of starting material were subjected to reduction/alkylation followed by on-bead digestion with trypsin. Protein identification using 1200 Easy-nanoLC - Orbitrap Exploris 480 platform (Thermo Fisher Scientific), typically generates up to 1000 protein identifications for similar samples, which being compared to control immunoprecipitated samples provided information on interacting partners. For each experimental group, 4 biological replicates were performed. Protein identification was done using Proteome Discoverer (Thermo Fisher Scientific) software, followed by analysis of up-regulated proteins using a 1% false discovery rate, identified by unique peptides.

### Bioinformatics analyses

The relative fluorescence unit (RFU) values for each protein from Mass Spectrometry were converted to Log2 values. The difference in Log2 values between rapamycin-treated and control samples was calculated to find the delta Log2 value. These delta Log2 values were further converted into fold-change values. Protein expression changes were analyzed statistically using a two-tailed Student’s T-test and Z-score, based on four protein expression replicates. Proteins showing significant expression changes (P-value < 0.05, Z-score ≥ +1.96σ or ≤ -1.96σ, and fold change > ±2) were chosen for further analysis with Ingenuity Pathway Analysis (IPA) software. IPA was used to predict the role of these proteins in cellular functions, signaling pathways, binding interactions, and to determine potential activation or inhibition of these pathways and functions, based on protein number and expression patterns.

### Preparing Trx1KD stable HT22 cells

HT22 cells were seeded in T75 flasks so that the cells become ∼60 - 70 % confluent on the next day. HT22 cells were transfected with psPAX2 (Addgene # 12260), and pMD2.G (Addgene # 12259) packaging vectors and with the transfer plasmid (Txn1 sgRNA CRISPR/Cas9, ABM Inc# 48866114) using lipofectamine 2000 (Invitrogen). Cells were cultured in the transfection media for 6 hours, then the media was changed with regular FBS DMEM growth media. After 24 hours, cells were seeded in 24 well plates with serial dilution and allowed to grow for 24 hours. HT22 cells were then treated with 8 µg/ml of puromycin (TOCRIS, 4089) in FBS/DMEM media. A single colony was picked and expanded in 4 µM of puromycin to get stable HT22 cells with Trx1KD.

### Western blot analysis

Western blotting was performed as described previously (Islam et al., 2021). Briefly, cells were scrapped following treatment, collected by centrifugation at 4000 rpm for 5 min, and washed twice with ice-cold PBS. Cell pellets were then lysed in NP-40 lysis buffer (50 mM Tris HCl pH 8, 150 mM NaCl, 5 mM EDTA, 1% NP-40) with 1X protease and phosphatase inhibitors and kept on ice for 40 mins. Cell suspensions were then subjected to sonication (3×5 s) on ice and centrifuged at 12, 500g for 15 min at 4°C to obtain the supernatant. Protein concentration in the whole cell lysate was measured using Pierce BCA Protein Assay Kit. An equal amount of proteins was resolved in SDS-PAGE at constant voltage and transferred to the PVDF membrane using the Trans-Blot® Turbo™ Transfer Buffer and System. After blocking for 1 h with 5% non-fat milk in Tris-buffered saline containing 0.2% Tween 20 (TBS-T), the membrane was probed with primary antibodies overnight at 4 °C, and HRP conjugated secondary antibody for 1 h at room temperature. To assess equal loading, membranes will be re-probed with β-actin or GAPDH antibody. Densitometry measurements were performed using ImageJ software.

### Immunocytochemistry

After the indicated treatments, cells were washed in PBS and fixed with 4% paraformaldehyde (PFA) in PBS for 15 min at room temperature or ice-cold 98% methanol for 10 min at -20 °C. Subsequently, cells were permeabilized with 0.3% Triton-X 100, and coverslips were incubated with appropriate primary antibodies overnight at 4°C followed by three 10 min washes with PBS. This was followed by a 2-hour incubation with the appropriate secondary antibody at room temperature. Coverslips were washed 3 times (10 min/wash) with PBS, followed by nuclear counterstaining with DAPI (5 min) and washed twice with PBS. Then, the coverslips were mounted on glass slides with mowiol. Immunofluorescence images were acquired by confocal microscope (Zeiss, LSM880) using similar settings across the experimental samples. All Alexa flour labelled secondary antibodies used for immunocytochemistry were purchased from Molecular Probes, USA. The following antibodies are used for this study: β-tubulin III (1:400, #), Actin phalloidin (1:400, Sigma Aldrich# A12379).

### F-Actin G - Actin Cytoskeleton Assay

Cellular F-actin and G-actin were determined by Triton X-100-insoluble cytoskeleton assay as described before [39]. In brief, primary cortical neurons and HT22 cells were lysed in a buffer - A containing 10 mM HEPES (pH 7.0), 1 mM MgCl2, 2.5 mM EGTA, 1% Triton X-100 and kept for 10 minutes at room temperature and centrifuged for 4 minutes at 8000 g. The supernatant containing G-actin was transferred to a new tube. F-actin containing pellet was washed twice with buffer A and solubilized with a buffer consisting of 0.1 M PIPES (pH 6.9) 10 mM CaCl2, 1 mM MgSO4 and 5 μM cytochalasin D and 6M Urea to depolymerize F-actin. Equal volumes of G-actin and F-actin fraction were separated on Sodium dodecyl sulfate-Polyacrylamide gel electrophoresis (SDS-PAGE) gels and western-blot based detection was performed using monoclonal anti-β-actin antibody (1:4000, #sc-47778, Santa Cruz Biotechnology, USA). Band signal intensity was detected by Clarity and Clarity Max ECL Western Blotting Substrates (BioRad) and F/G-actin ratio was measured by densitometry of immunoblots.

### Quantification of F-actin in immunocytochemistry

F-actin puncta were quantified in mouse primary neurons as described previously (H. Li, Aksenova, Bertrand, Mactutus, & Booze, 2016). In brief, three coverslips of mouse cortical neurons per condition was immunolabeled with actin-phalloidin and β-tubulin III antibodies. Images of co-labeled F-actin/β-tubulin III neurons were acquired under far red (647 nm)/orange (568) fluorescent channels. Five fluorescent images of individual neurons with clearly defined dendritic arbors were chosen for quantification. F-actin rich structures in second order dendritic segment (length range 25-75 μm) with continuous β-tubulin III signal was identified. After subtraction of the background, bright red F-actin puncta were counted (size ≤1.5 µm) along the dendrite length. The length of selected dendritic segment was determined, and the density was calculated by dividing total F-actin labeled puncta (N) by the length (L) of the β-tubulin III labeled dendrites. Data was expressed as number of F-actin puncta per μm of dendrite. Quantification of actin fiber lengths was done using Ridge Detection plugin in ImageJ [40, 41] with lower threshold 3.00, upper threshold 8.00 and sigma 1.50.

### Trx1 Knocked down in rat primary neuronal cultures

Dissociated cortical neuronal cultures were prepared from rat embryos at day 18 (E-18) as described before [42]. 300,000 cells per 60 mm culture dish were seeded on poly-L-lysine-coated coverslips and inverted over a glial feeder layer. Two days after plating, cytosine arabinoside (5 mM) was added per dish of neuron culture to prevent glial cell outgrowth. We employed genetic manipulation of neurons by performing transfection of neurons with scramble or Trx1 short hairpin RNAs (shRNAs). Transfection was done by nucleofection (AMAXA Biosystems, Lonza) with 4-5 ug DNA at day DIV 0 and seeded at a density of 1.5 million per 60 mm dish.

### Statistical Analysis

Graph Pad Prism® version 6 was used for statistical analyses. The data are reported as mean ± SEM. To compare between more than two groups, the statistical analysis was carried out by one-way analysis of variance (ANOVA) followed by Tukey’s post-hoc test. To compare between two groups, a two-tailed unpaired student t-test was used. p ≥ 0.05 were considered as statistically significant.

## Acknowledgements

EE is supported by a Discovery Grant from Natural Sciences and Engineering Research Council of Canada (RGPIN-2019-05371). This work was partially supported by the Canadian Institute of Health Research Project Grant: 156218 to S. K-A.

